# Induction of senescence upon loss of the Ash2l core subunit of H3K4 methyltransferase complexes

**DOI:** 10.1101/2022.02.11.480149

**Authors:** Agnieszka Bochyńska, Alexander T. Stenzel, Roksaneh Sayadi Boroujeni, Chao-Chung Kuo, Mirna Barsoum, Weili Liang, Philip Bussmann, Ivan G. Costa, Juliane Lüscher-Firzlaff, Bernhard Lüscher

## Abstract

Gene expression is controlled in part by post-translational modifications of core histones. Methylation of lysine 4 of histone H3 (H3K4), associated with open chromatin and gene transcription, is catalyzed by type 2 lysine methyltransferase complexes that require WDR5, RBBP5, ASH2L and DPY30 as core subunits. Ash2l is essential during embryogenesis and for maintaining adult tissues. To expand on the mechanistic understanding of Ash2l, we generated mouse embryo fibroblasts (MEFs) with conditional *Ash2l* alleles. Upon loss of Ash2l, methylation of H3K4 and gene expression were downregulated, which correlated with inhibition of proliferation and cell cycle progression. Moreover, we observed induction of senescence concomitant with a set of downregulated signature genes but independent of SASP. Many of the signature genes are FoxM1 responsive. Indeed, exogenous FOXM1 was sufficient to delay senescence. Thus, although the loss of Ash2l in MEFs has broad and complex consequences, a distinct set of downregulated genes promotes senescence.

## INTRODUCTION

The regulated expression of defined genes is important for the identity and physiology of cells. Transcription of genes is regulated by sequence specific transcription factors and by cofactors that control chromatin accessibility and polymerase activity. The smallest units of chromatin are the nucleosomes, which are composed of 8 core histones, two copies each of H2A, H2B, H3 and H4 or various variants thereof. Several hundred post-translational modifications (PTM) of core histones have been described, which are thought to regulate specific aspects of transcription ^1, 2, 3^. Methylation of histone H3 at lysine 4 has gained attention because its mono- and tri-methylation (H3K4me1/3) is associated with active enhancers and promoters, respectively. This PTM is catalyzed primarily by lysine-specific methyltransferases of the KMT2 family, consisting of MLL1 to 4, and SET1A and B (KMT2A-D, F and G, respectively). These enzymes are part of multi-subunit complexes, referred to as COMPASS (complex of proteins associated with Set1). All six KMT2 family members associate with the WRAD core complex composed of WDR5, RBBP5, ASH2L, and two copies of DPY30, forming functional KMT2 complexes. WRAD is necessary for catalytic activity of KMT2 enzymes. Additional subunits interact specifically with distinct KMT2 complexes, further increasing the diversity and functionality of these methyltransferases ^4, 5, 6, 7, 8, 9^.

The importance of KMT2 complexes is highlighted by many functional studies. All KMT2 complex subunits, which have been knocked-out, are essential in mice ^7, 8^. This is true for the 6 enzymatic subunits, which are required during defined developmental processes, indicating at least in part distinct functions. Moreover, the two WRAD core complex subunits, Ash2l and Dpy30, are necessary for organismal development and cell proliferation and differentiation in the mouse ^10, 11, 12, 13, 14^. Of note is also that mutations in the *Drosophila melanogaster* orthologue *Ash2* cause homeotic transformations, similar to mutations in *trithorax*, the orthologue of *MLL1* and *MLL2* ^15, 16, 17^. Moreover, KMT2 complex subunits are linked to various diseases, including cancer ^7, 18^. Together, KMT2 complexes/COMPASS contribute to the control of cell proliferation and differentiation and participate in cell fate decisions.

We identified ASH2L as an interaction partner of the oncoprotein c-MYC and our findings suggest that ASH2L has oncogenic activity ^19, 20^. In mice, the loss of Ash2l led to disintegration of hepatocytes ^14^. In hematopoietic stem and progenitor cells (HSPCs), the knockout of *Ash2l* resulted in an accumulation of cells in a G2/M cell cycle arrest, accompanied with a proliferation and differentiation stop, that resulted in a loss of mature cells ^13, 14^. We have now addressed the function of Ash2l in mouse embryo fibroblasts (MEFs). Unlike the findings in hepatocytes and HSPCs, Ash2l loss in MEFs resulted in a senescence phenotype once cells ceased to proliferate. Senescence is a heterogenous response of cells to a broad spectrum of triggers, which include replicative exhaustion with telomer erosion, oncogene activation, and DNA damage as well as other forms of stress that result in the upregulation of many genes, including those expressing cyclin-dependent kinase inhibitors ^21, 22^. Ash2l loss was accompanied by an overall reduction in gene expression. A small set of downregulated genes are linked to senescence that are also repressed in other senescent cells. Many of these genes are controlled by the transcription factor FoxM1. Its exogenous expression was sufficient to delay senescence. Thus, MEF cells respond to Ash2l loss with a senescence program that depends on downregulation of genes associated with the cell cycle, replication and DNA repair, which is in contrast to the typically observed upregulation of genes.

## RESULTS

### Ash2l is required for fibroblast cell proliferation

We prepared MEF cells from mouse embryos that contained a floxed *Ash2l* exon 4 and that express a Cre-ER^TM2^ recombinase fusion protein. Primary MEFs (pMEFs), were immortalized using an siRNA expression construct that targets the *p19*^*ARF*^ mRNA (iMEFs) ^23, 24^. Two pairs of iMEF cells were generated, WT1 and WT2 cells (*Ash2l*^*wt/wt*^:*Cre-ER*^*TM2*^) and KO1 and KO2 cells (*Ash2l*^*fl/fl*^:*Cre-ER*^*TM2*^), each pair from 2 embryos of the same mother. Upon treatment with 4-hydroxy tamoxifen (HOT), which activates the constitutively expressed recombinase, exon 4 sequences were lost in KO but not in WT cells (Suppl. Fig. S1A) and exon 4 containing RNA expression decreased strongly (Fig. 1A). This was followed by a decrease of Ash2l in KO2 cells (Fig. 1B) and in KO1 and pMEF cells (Suppl. Fig. S1B and C), albeit slowly due to the long half-life of Ash2l protein ^20^. As a result, H3K4 methylation decreased (Fig. 1C and Suppl. Fig. S1B-D), as expected when one of the core components of KMT2 complexes is lost. Moreover, Rbbp5 containing complexes lost H3K4 methyltransferase activity when isolated from cells without Ash2l (Fig. 1D). Rbbp5, a direct interaction partner of Ash2l, was reduced at late time points, while Wdr5 was not (Fig. 1 and Suppl. Fig. S1). The loss of Ash2l did not affect RNA expression of the core components *Wdr5, Rbbp5*, and *Dpy30*, and the different catalytic subunits (Suppl. Fig. S1E). Thus, loss of Ash2l prevented H3K4 methylation, consistent with previous studies addressing the role of KMT2 complexes ^7, 8^.

**Figure 1.**
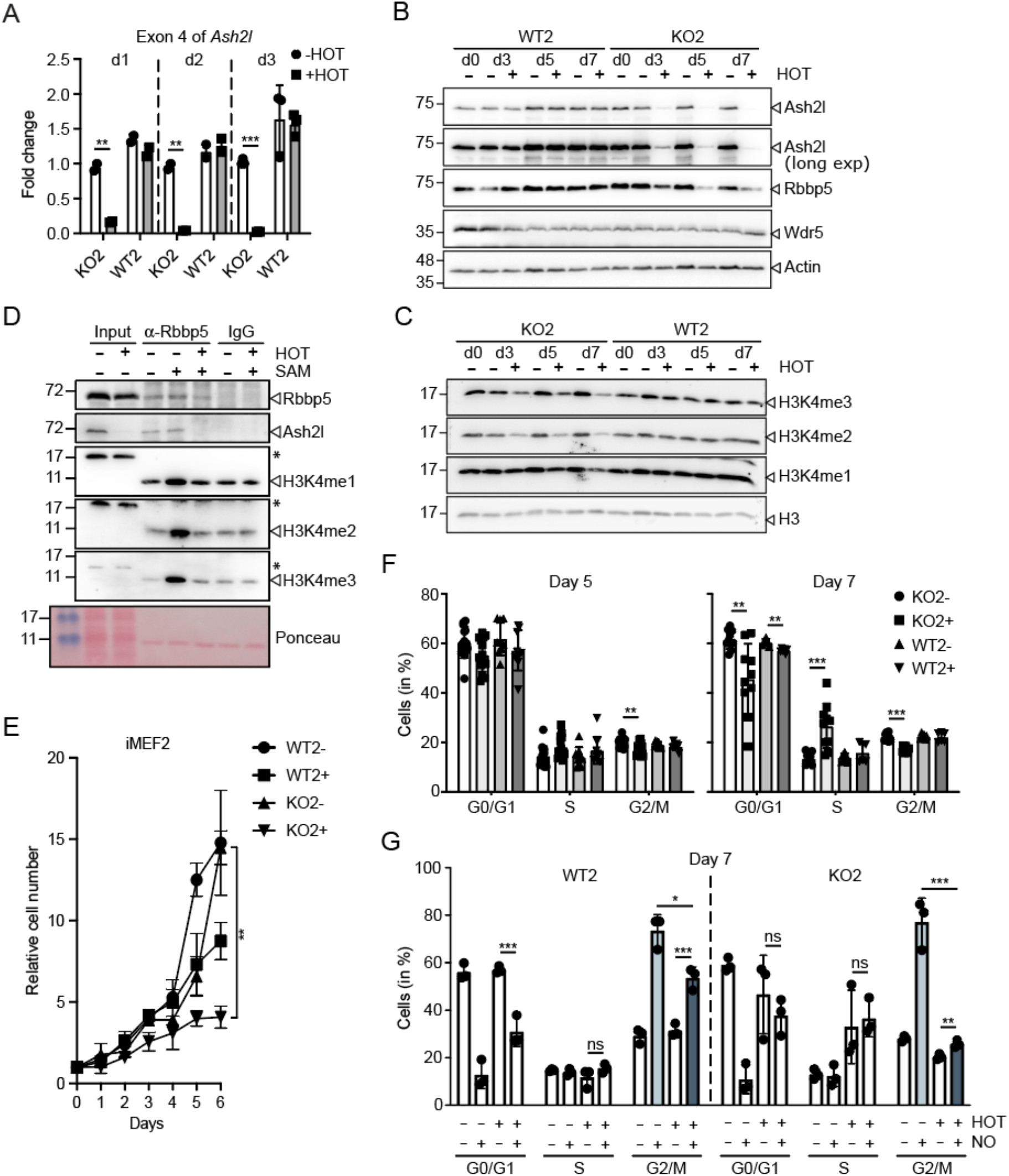
Loss of Ash2l in mouse embryo fibroblasts inhibits proliferation and cell cycle progression. A. iMEF2 cells (WT2, *Ash2l*^*wt/wt*^:*Cre-ER*^*TM2*^; and KO2, *Ash2l*^*fl/fl*^:*Cre-ER*^*TM2*^) were treated with 4-hydroxy tamoxifen (HOT, 5 nM) or vehicle as indicated. RT-qPCR analysis of the floxed exon 4 of *Ash2l* was performed at days 1, 2 and 3. Indicated are mean values ± SD (*** p<0.001; ** p<0.01). B and C. Wild type (WT1) and *Ash2l*^fl/fl^ (KO1) immortalized fibroblasts (iMEF1) were treated ± HOT for 0, 3, 5 or 7 days. The cells were lysed and the indicated proteins analyzed by Western blotting. D. KO2 cells were treated ± HOT for 5 days. KMT2 complexes were immunoprecipitated from low stringency lysates using an antibody specific for Rbbp5 or a species matched control antibody (IgG). The immunoprecipitates were incubated in the presence or absence of S-adenosyl-methionine (SAM) and recombinant histone H3. The reactions were analyzed for the proteins indicated using Western blotting. In the input non-specific signals are labeled (*). E. Wild type (WT) and *Ash2l*^fl/fl^ (KO) immortalized fibroblasts (iMEF2) were treated ± HOT (5 nM) for the times indicated. The cells were counted daily. Mean values ± SD of 4-6 measurements in duplicates for each time point (statistical analyses refer to day 6: ** < 0.01). F. Cell cycle analysis using flow cytometry of fixed and Hoechst stained WT2 and KO2 cells treated ± HOT for 5 or 7 days. Mean values ± SD (n = 6-10; ** < 0.01, *** < 0.001). G. WT2 and KO2 cells were treated ± HOT for 7 days. During the last 18 h the cells were incubated with nocodazole (100 ng/ml) or vehicle control as indicated. The cells were fixed, the DNA stained using Hoechst and analyzed by flow cytometry. Mean values ± SD of 3 experiments are shown. Statistical analyses are indicated for selected data sets, * <0.05, ** < 0.01, *** < 0.001, ns, not significant.

Ash2l loss inhibited MEF proliferation beginning between day 3 and 5 upon HOT treatment and ceased by day 6 (Fig. 1E and Suppl. Fig. S2A). This correlated with the decrease in Ash2l and in H3K4me3 and me1. Unlike in HSPCs, we did not observe an accumulation in the G2/M phase, rather the distribution of the cells in the different cell cycle phases remained largely unaffected with a small increase in S phase at day 7 (Fig. 1F and Suppl. Fig. S2B and C). This suggested that the cells were unable to transit the cell cycle upon downregulation of Ash2l. To address whether the cells were stalled in the cell cycle, proliferating iMEFs were treated for 18 hours with nocodazole, which arrests cells in early mitosis ^25^. In the absence of HOT, both WT2 and KO2 cells accumulated in G2/M, concomitant with a decrease in G0/G1 cells (Fig. 1G). Similarly, an increase in G2/M phase, albeit smaller, was seen in HOT treated WT2 cells. WT cells proliferated slower upon HOT treatment (Fig. 1E and Suppl. Fig. S2A), explaining the reduced G2/M accumulation. In the absence of Ash2l, the accumulation of KO2 cells in G2/M was minimal (Fig. 1G), indicating that these cells were unable to transit through the cell cycle.

We also considered that the loss of Ash2l induced apoptosis, thereby influencing overall proliferation. We did not notice an increase in sub-G0/G1 cells in flow cytometry (Suppl. Fig. S2C). Consistent with this observation, staining cells for annexin V and with propidium iodide (PI) did not reveal significant changes, neither in early (annexin V positive) nor late apoptotic cells (both annexin V and PI positive) (Suppl. Fig. S2D and E). Thus, cell death is unlikely to contribute significantly to the reduced cell proliferation.

### Loss of Ash2l promotes senescence

We noticed that the cells became larger upon loss of Ash2l, an attribute of senescent cells (Fig. 2A). Indeed, many cells became SA-β-galactosidase (SA-β-gal) positive (Fig. 2A und B), a frequently used marker for senescence ^22^. This phenotype was further assessed by applying senolytic drugs. These have been described to selectively kill senescent cells ^26^. For example, the combination of Dasatinib and Quercetin eliminated senescent MEFs ^27^. We observed a reduction of more than 50% of SA-β-gal positive cells with 250 nM Dasatinib/15 µM Quercetin and a trend to less senescent cells with 25 nM Dasatinib/15 µM Quercetin (Fig. 2C), while 10 µM Dasatinib/15 µM Quercetin, the originally described dose for MEFs ^27^, efficiently killed all our cells. Moreover, it was noted that the knockdown of phosphoinositide 3 kinase delta (PI3Kδ) resulted in a senolytic effect ^28^. Inhibiting PI3 kinases with Wortmannin or compound 15e reduced the number of senescent cells (Fig. 2C) ^29, 30^. One of the signaling pathways that is activated during senescence involves p38MAPK, a stress response kinase ^31^. Indeed, the loss of Ash2l resulted in activating phosphorylation of p38MAPK (Fig. 2D). Thus, the loss of Ash2l in MEF cells promotes senescence.

**Figure 2.**
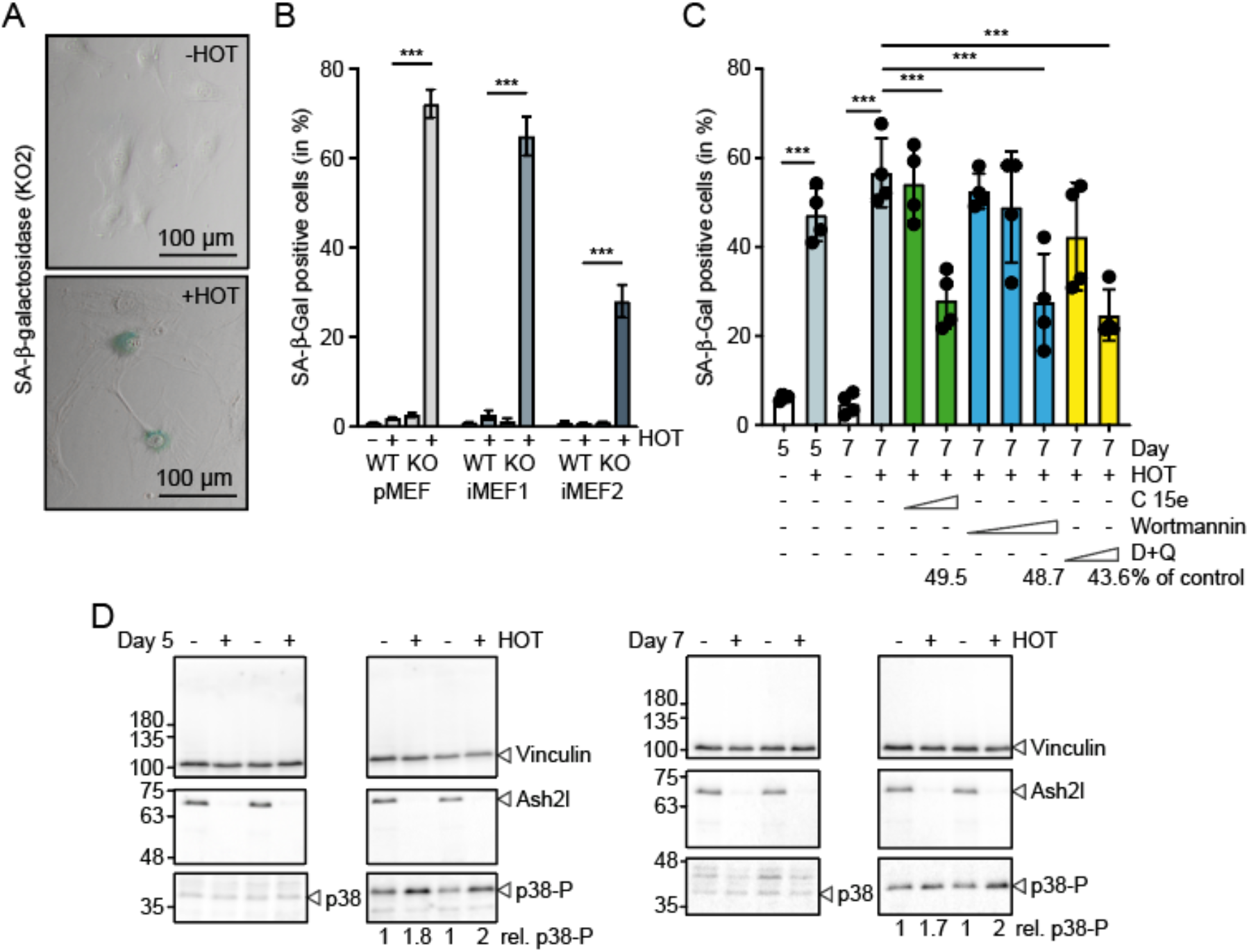
Ash2l loss promotes senescence in MEF cells. A. KO2 cells were incubated ± HOT for 7 days. The cells were fixed and stained for SA-β-galactosidase activity (blue). B. Primary MEF (pMEF) and iMEF cells were treated ± HOT for 5 or 7 days as indicated, analyzed as in panel A and the number of SA-β-gal positive cells determined. Mean values ± SD of 3-4 experiments are shown. In each experiment 200-300 cells were counted (*** < 0.001). C. KO2 cells were incubated ± HOT for 7 days. During the last 2 days senolytic drugs were added: Compound 15e (1 and 5 µM), Wortmannin (0.1, 1 and 10 µM), and a combination of Dasatinib/Quercetin (25 nM/1.5 µM and 250 nM/15 µM). The percentage of SA-β-Gal positive cells is displayed. Four sections each with 50-150 cells of two biological replicates were counted. Mean values ± SD (n = 4; *** < 0.001). D. KO2 cells were treated ± HOT for 5 or 7 days (top and bottom panels, respectively). Whole cell lysates of two biological replicates were analyzed by Western blotting using the indicated antibodies. Quantified were p38-P vs. p38 signals.

### Downregulation of gene expression upon loss of Ash2l

A hallmark of senescent cells is the secretion of signaling and extracellular matrix modulating molecules, at least in part due to an altered gene expression program, referred to as senescence-associated secretory phenotype (SASP) ^22, 32, 33^. Because KMT2 complexes are associated with altered gene transcription, we performed RNA-seq experiments to evaluate changes in gene expression, which might account for the senescence phenotype, upon Ash2l loss five days after HOT addition (Suppl. Table S1). In view of the strong correlation of H3K4me1 and me3 with active enhancer and promoters, respectively, we expected broad effects on gene expression and cellular RNA content. To compensate for this, we used ERCC (External RNA Controls Consortium) spike-in RNA. Roughly 75% of all genes (37205) showed reduced expression (Log_2_FC < 0) in the HOT treated KO1 and KO2 cells (Fig. 3A and B). We observed 1118 and 1600 significantly downregulated genes in KO1 and KO2 cells, respectively (Fig. 3A and C). Also, a small number of genes were up-regulated (176 and 207 in KO1 and KO2 cells, respectively) (Fig. 3A and C). Of note, all the apparently upregulated genes had very low basal expression (Suppl. Table S1).

**Figure 3.**
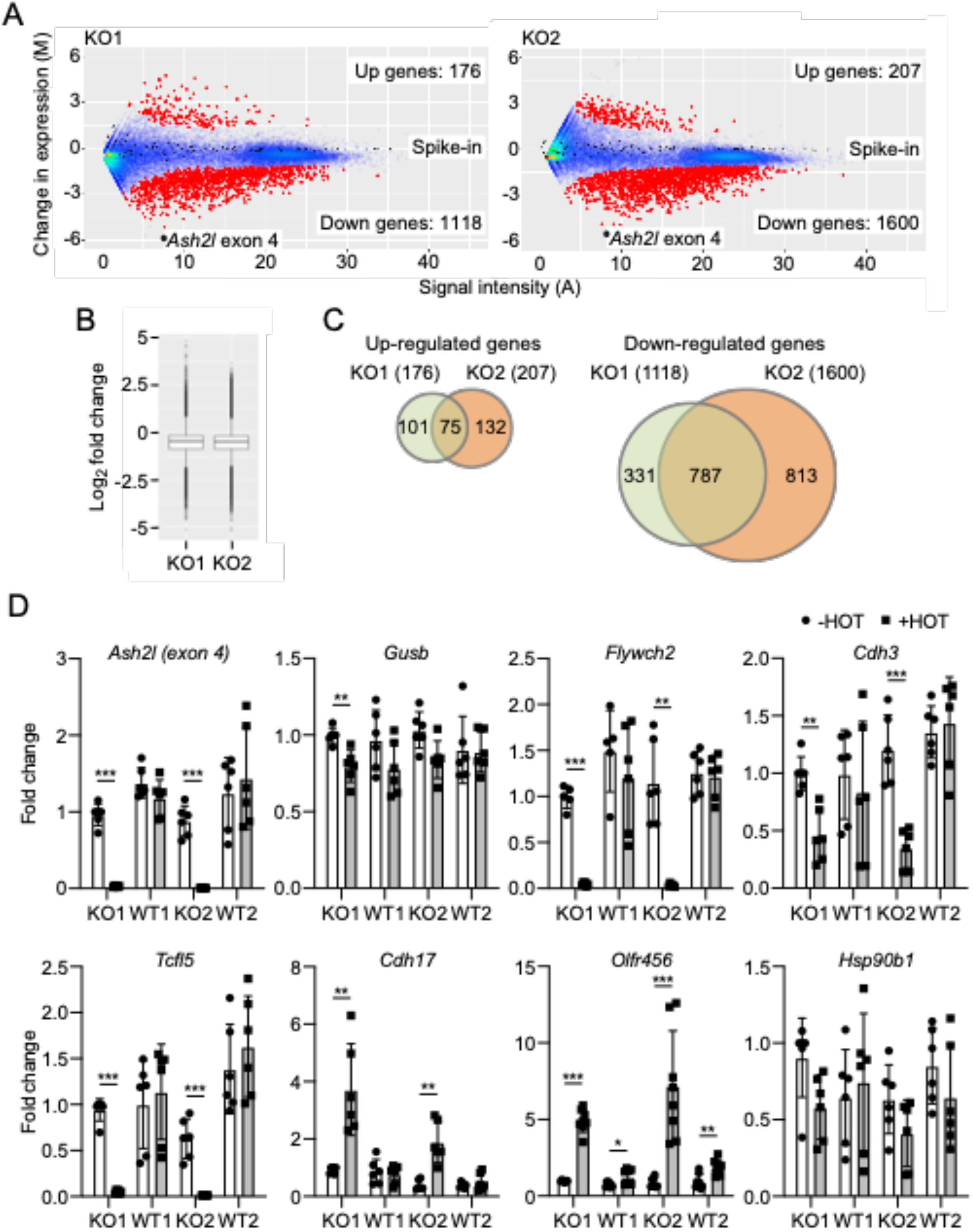
Deregulated gene expression upon loss of Ash2l. A. KO1 and KO2 cells were treated ± HOT for 5 days. RNA was isolated and ERCC RNA spike-in control mix added. The RNA was analyzed using next generation sequencing, adjusted to the spike-in RNA. Displayed are MA plots (A = 0.5(log_2_(KO+HOT) + log_2_(KO-HOT)); M = log_2_(KO+HOT) – log_2_(KO-HOT)). The number of up- and downregulated genes are indicated (p-value < 0.05). Red dots, significantly regulated genes; blue dots, not regulated genes; black dots, spike-in RNA. The signals obtained for exon 4 of *Ash2l* are indicated. B. Box plot analysis of gene expression in KO1 and KO2 cells from the data shown in panel A. A one sample t-test indicates that FC values deviate from 0 (p-value < 2.2e- 16). C. Comparison of up- and downregulated genes between KO1 and KO2 cells. D. WT and KO iMEF1 and 2 cells were incubated ± HOT for 5 days, the RNA isolated and the expression of the indicated genes measured by RT-qPCR. The expression of *Gusb* was used as reference. Mean values ± SD (n = 5-8) (* <0.05, ** < 0.01, *** < 0.001).

Altered gene expression was verified in independent RT-qPCR experiments. *Ash2l* exon 4 expression was strongly decreased as expected (Fig. 3A and D). The expression of *β-glucuronidase (Gusb)* was only slightly downregulated, consistent with the trend to decreased expression of many genes, and served as control (Fig. 3D). RNA-seq revealed that *Flywch2, Tcfl5* and *Cdh3* were downregulated, the former two being in the top group. *Cdh17* and *Olfr456* were up-regulated while the expression of *Hsp90b1* showed little change (Suppl. Table S1). The differential expression of these genes was reproduced in the RT-qPCR experiments (Fig. 3D). Thus, the loss of Ash2l resulted in broadly deregulated gene expression.

We employed gene ontology (GO) analyses to evaluate whether the deregulated genes would allow to make predictions about the cellular processes that might be affected upon Ash2l loss, particularly considering a link to senescence. A large number of terms were observed for downregulated genes (Suppl. Fig. S3A), When up- regulated genes were analyzed, only 4 GO terms were statistically significant (Suppl. Fig. S3B). The terms were very broad, but included some related to cell cycle and proliferation. Nevertheless, terms that would suggest a direct link to senescence were not observed in either group.

### Gene repression is associated with senescence

One of the key events is the induction of SASP that arguably occurs in all senescent cells ^22^. Because SASP is primarily based on gene induction, it is unlikely that this can be observed in the *Ash2l* KO cells as the major response is gene repression (Fig. 3). Therefore, we compared our downregulated genes with those of two studies ^34, 35^. In one senescence was induced in three primary human cell populations, i.e. umbilical vein endothelial cells, fetal lung fibroblasts, and mesenchymal stromal cells, by replicative exhaustion ^35^. Gene expression was measured when roughly 65% senescent cells had accumulated as determined by SA-β-gal staining. Of the several hundred downregulated genes in each cell population, 206 were in common between the 3 cell populations ^35^. We found that 37 and 87 homologous genes were significantly downregulated in KO1 and KO2 cells, respectively, despite comparing human and mouse expression patterns. Thirty-three of these genes were downregulated in both KO iMEF populations (Fig. 4A and B, B+M genes). In the second study, replicative senescence was analyzed in human diploid fibroblasts ^34^. Downregulated genes were enriched for pathways associated with proliferation and replication. We observed an overlap of 72 genes with our dataset (Fig. 4A and B, A+M genes). Moreover, the genes in the M group are also downregulated in other studies of cellular senescence ^36, 37^. Importantly, GO pathway analyses of these genes (A, B and M) revealed terms associated with cellular senescence (Fig. 4C, e.g. cellular senescence, p53 signaling pathway). Consistent with inhibition of proliferation and cell cycle progression were additional terms (Fig. 4C, e.g. Faconi anemia pathway, p53 signaling pathway, FoxO signaling pathway) and the repression of many genes associated with the cell cycle and replication (Suppl. Table S2). Also the terms relating to virus infection are consistent with a proliferation stop as cellular response and senescence (Fig. 4C) ^38^. Together, these changes in gene expression render further support for induction of senescence upon Ash2l loss.

**Figure 4.**
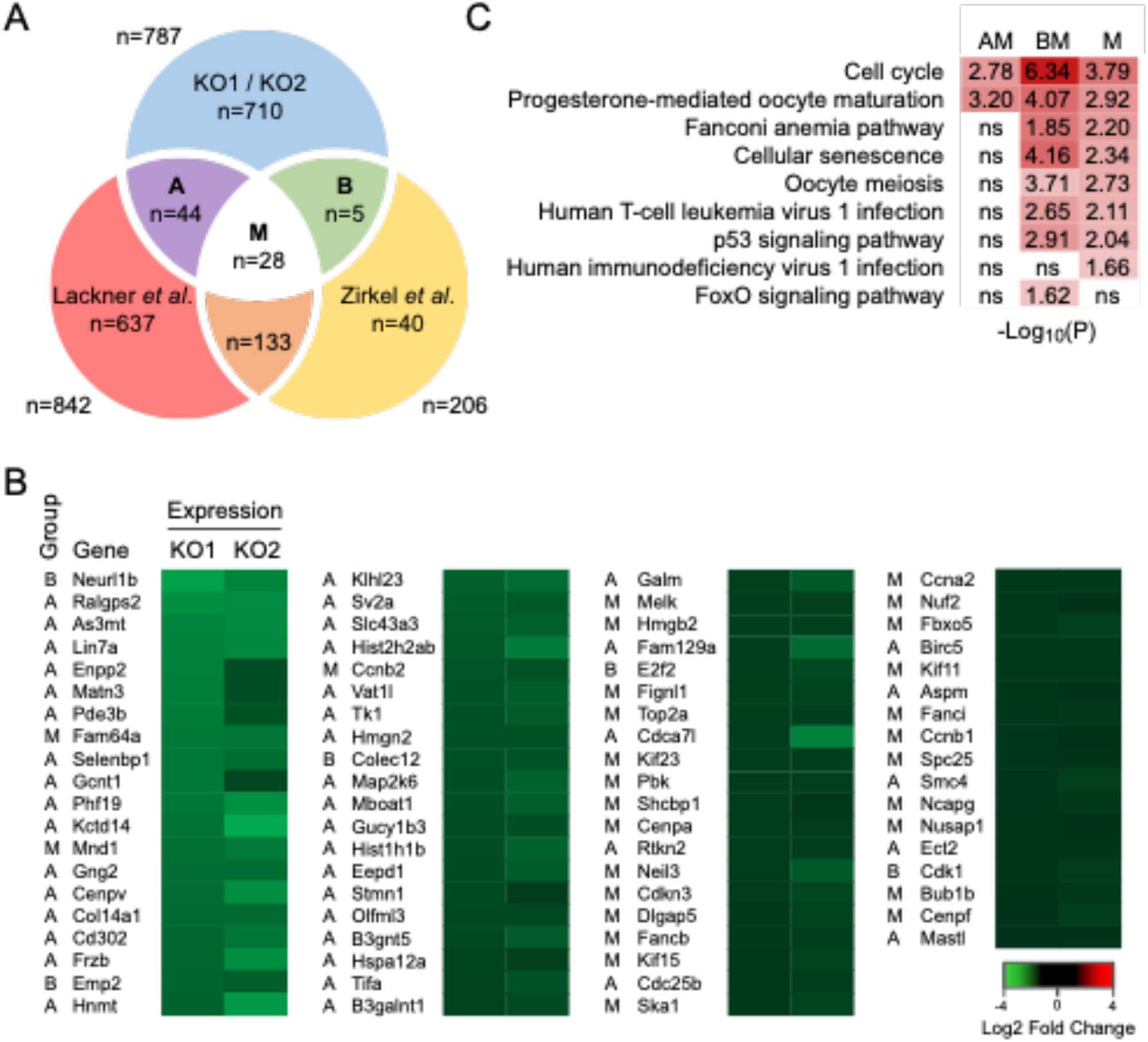
A. Commonly downregulated genes in murine KO1 and KO2 cells upon loss of Ash2l were compared to commonly downregulated genes in 3 primary human cell populations ^35^ and to downregulated genes in human diploid fibroblasts ^34^ in response to replicative senescence. The number of genes is indicated that are in common between the different groups: A, KO1/KO2 and the genes in Lackner et al.; B, KO1/KO2 and the genes in Zirkel et al.; M, common genes of all three studies. Comparison KO1/KO2 with Lackner et al. and Zirkel et al.: p-value < 2.2e-16 (Fischer Exact test). B. The genes in groups A, B and M are listed and the repression of these genes in both KO1 and KO2 cells upon treatment ± HOT for 5 days is indicated. C. GO pathway analysis of the genes that are in common between the data sets of panel D. Adjusted p-values are indicated; ns, not significant.

To obtain information about the dynamics of events, e.g. whether a transient SASP response might occur, we performed time course experiments. The first SA-β-gal positive cells appeared at day 4 (Fig. 5A), concomitant with inhibition of cell proliferation (Fig. 1E and S2A). The loss of Ash2l and the decrease in H3K4 methylation may affect chromatin and provoke transcription-replication conflicts ^39^, which may result in DNA damage that is known to contribute to induction of senescence ^40^. Therefore, we measured γ-H2AX, the phosphorylated form of histone H2AX, a marker for double strand DNA breaks ^41, 42^. Occasionally we observed Ash2l negative cells that stained positive for γ-H2AX at day 5 (Suppl. Fig. S4A). However, no broad induction of γ-H2AX in response to Ash2l loss was observed prior to significant senescence induction at days 5 and 7 (Suppl. Fig. S4A).

**Figure 5.**
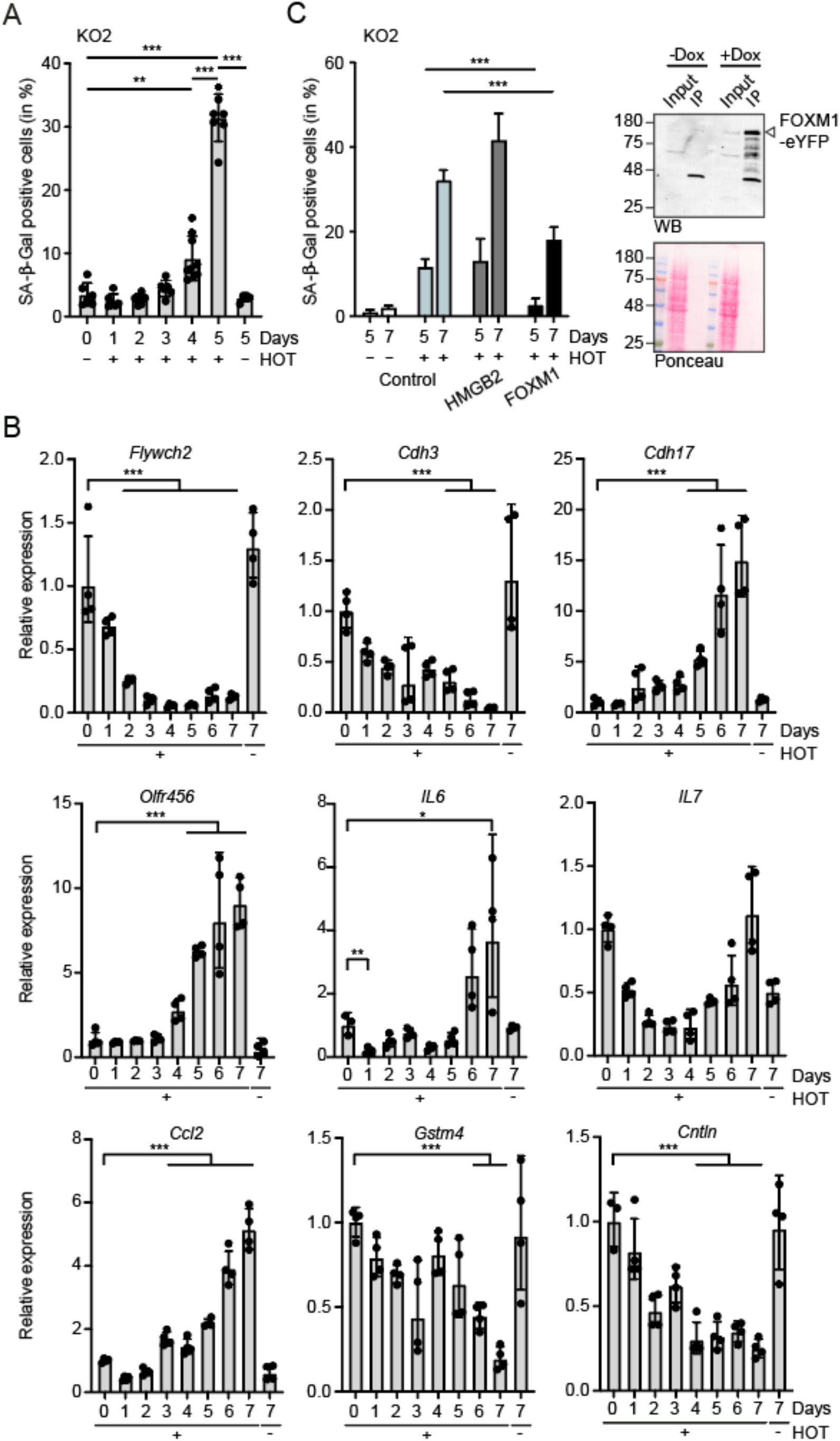
FOXM1 delays Ash2l loss induced senescence. A. KO2 cells were treated with ± HOT for the indicated days. Then cells were fixed and stained for SA-β-galactosidase activity and counted. Mean values ± SD (n = 6-7) (p = * <0.05, ** < 0.01, *** < 0.001). B. KO2 cells were incubated ± HOT for 1-7 days, the RNA isolated and the expression of the indicated genes measured by RT-qPCR. The expression of *Gusb* was used as control. Mean values ± SD (n = 4) (* <0.05, ** < 0.01, *** < 0.001). C. KO2 cells were infected with lentiviruses expressing HMGB2 and FOXM1 (as eYFP fusion proteins). Pools of cells were treated with HOT as indicated and with doxycycline (Dox). The number of SA-β-galactosidase positive cells was determined at days 5 and 7. Slides of two biological replicates analyzed in duplicates were blinded and counted by 3 persons. The resulting mean values ± SD (*** < 0.001) are depicted. The panel on the right documents FOXM1-eYFP expression in KO2 cells upon induction with Dox using Western blotting (WB). Input refers to 2.5% of total cell lysate, the remainder was used for immunoprecipitating the fusion protein (IP). Prior to specific protein detection, the membranes were stained with Ponceau S to verify equal loading of input lysates.

Because SASP is a prominent feature of senescent cells, we considered that genes expressing SASP factors might be induced transiently, prior to the appearance of SA- β-gal positive cells. This might occur prior to inhibition of cell proliferation, which we may have missed in our RNA-seq experiments. Therefore, we measured RNA expression over 7 days upon HOT treatment. For control, two downregulated, *Flywch2* and *Cdh3*, and two up-regulated genes, *Cdh17* and *Olfr456*, were measured (Fig. 5B). *Flywch2* and *Cdh3* were strongly repressed by day 3 while induction of RNA expression was a late event. This supports the notion that upregulation of genes was most likely an indirect consequence of Ash2l loss. We analyzed three SASP genes, *IL6, IL7* and *Ccl2*, that show basal expression in MEF cells and which are typically upregulated in senescent cells ^22^. All three were initially downregulated. At later time points, parallel to the appearance of SA-β-gal positive cells, *IL6* and *IL7* expression was enhanced (Fig. 5B). At present it is unclear how the mRNAs of these genes are upregulated. Additionally, we analyzed two genes, *Gstm4* and *Cntln*, that were identified as downregulated core senescence signature genes in three different cell types, including human foreskin fibroblasts ^43^. Both were also repressed in our iMEF cells (Fig. 5B).

### The transcription factor FOXM1 delays senescence induced by Ash2l loss

When the M genes were analyzed, it was obvious that many fulfill functions in the regulation of the cell cycle, in particular the metaphase – anaphase transition, and in DNA repair and genomic stability, consistent with the GO analysis (Fig. 4C). Many of the genes in the M group are regulated by the transcription factor FoxM1 (e.g. ^44, 45, 46, 47,48^), some of which antagonize senescence (e.g. ^49, 50, 51^). Moreover, FoxM1 itself is promoting cell cycle progression, having functional relevance both at the G1-S and the G2-M transitions (e.g. ^52, 53, 54, 55, 56^). Also, in several cellular systems, the exogenous expression of FoxM1 was sufficient to interfere with induction of senescence (e.g. ^57, 58, 59, 60, 61^). High mobility group proteins, including Hmgb2, possess functions in the control of chromatin and its downregulation is an early marker for senescence and aging ^35, 62, 63^. Conversely, the expression of HMGB2 is induced when cells enter the cell cycle from a quiescent state ^64^. A role of HMGB2 in senescence is further supported by its ability to inhibit the spreading of heterochromatin, which ensures expression of SASP genes ^65^. Based on these published findings, we generated iMEF cells in which we introduced inducible expression constructs for FOXM1 and HMGB2, both fused to eYFP, into KO2 cells. Pools of cells were then treated with HOT and the induction of the transgenes induced with doxycycline. The expression of FOXM1 delayed the development of senescent cells, suggesting that FOXM1 and its downstream targets are relevant downregulated genes in the Ash2l model (Fig. 5C). Unlike FOXM1, HMGB2, was not affecting senescence, despite its potential broad role in regulating chromatin and gene transcription.

## Discussion

We report on the consequences of the loss of Ash2l in MEF cells. Ash2l is necessary for proliferation and cell cycle progression of these cells. The loss of Ash2l does not result in a specific block in the cell cycle but rather cells appear to arrest through all phases. Moreover, we did not observe any cell death, rather the cells developed a flat, senescence-like phenotype. Consistent with senescence, iMEF cells became SA-β-gal positive. This appears to be the consequence of broad downregulation of gene expression. In particular we noticed a set of downregulated genes that are also apparent in other senescent cells. However, no SASP was observed, which is a hallmark broadly detectable in senescent cells. We noticed that many of the downregulated genes, which are linked to cell cycle progression and stress response, are controlled by the transcription factor FOXM1. This factor is important for cell cycle progression, particularly at the G1-S and G2-M transitions. When *Ash2l* was deleted in cells that express exogenous FOXM1, senescence was delayed. Together the findings suggest that downregulation of FOXM1 target genes is part of the circuitry that promotes senescence upon Ash2l loss and broad downregulation of gene transcription.

The loss of Ash2l results in depletion of H3K4 methylation, consistent with the observations that the WRAD core complex is necessary for KMT2 catalytic activity (Fig. 1) ^66, 67, 68, 69, 70, 71^. H3K4me1 and H3K4me3 correlate well with active enhancers and promoters, respectively ^4, 8, 9^. This is coherent with the observation that most genes are downregulated (Fig. 3). A number of suggestions have been made how H3K4 methylation might affect gene transcription. H3K4me3 provides binding sites for cofactors and thus may contribute to the local organization of chromatin ^72, 73, 74, 75, 76^. One of the readers that binds to H3K4me3 is TAF3, a subunit of the TFIID complex and thus intimately associated with Pol II loading onto core promoters ^77^. TAF3 possesses a PHD finger that recognizes H3K4me3 ^78^. This interaction is antagonized by asymmetric demethylation of H3 arginine 2 (H3R2me2a), which also affects methylation of neighboring K4 ^79^. Localizing H3R2me2a suggests that it is excluded from active promoters and thus provides evidence for crosstalk between these two marks. The essential nucleosome remodeling factor NURF can also bind to H3K4me3 and thus may provide local, promoter associated chromatin remodeling activity ^80, 81^. Multiple additional factors, including inhibitor of growth (ING) proteins and the chromatin associated factor PHF13/SPOC1, were identified as readers of H3K4me3 that might be associated with gene transcription ^5, 82, 83^. Thus, the loss of H3K4 methylation at promoters is likely to have multiple consequences, many are consistent with reduced gene expression.

In hematopoietic cells, the loss of Ash2l resulted in a cell cycle arrest in late G2/early M phase and as a consequence differentiation to mature cells was inhibited ^13^. Similarly, Dpy30 is necessary for hematopoietic cell proliferation and differentiation. *Dpy30* KO hematopoietic cells also accumulate at the HSPC state ^12, 84^, suggesting that the main phenotype observed upon loss of either Ash2l or Dpy30 is due to the inactivation of KMT2 complexes. However, unlike in HSPCs, the fibroblasts did not accumulate in any particular cell cycle phase, with the exception of a small increase in S phase cells (Figs. 1). Among the downregulated genes were many that encode cell cycle regulators, such as different cyclins, Cdk1 and E2F transcription factors ^85^, and replication factors, including 4 out of 6 minichromosome maintenance proteins (Mcm) (Table S2). The Mcm2-7 complex functions together with additional proteins as essential helicase during DNA double strand opening ^86^. Thus, the observed cell cycle phenotype is in agreement with the repression of genes that encode essential regulators of both cell cycle progression and replication. We assume that depending on the process that becomes limiting first, cells arrest in different phases of the cell cycle.

The phenotypic outcome of Ash2l loss in iMEF cells is senescence. This is a highly heterogeneous process when comparing different cell types and conditions. Stress-induced irreversible proliferation arrest and resistance to both mitogenic and oncogenic stimuli have been suggested to best define the senescent state ^22, 33, 40^. Also the observed cell cycle arrest and the lack of apoptosis are features of senescent cells ^87^. Typical stressors include excessive replication (replication stress), the induction of oncogenes (oncogene-induced senescence), and DNA damage (e.g. ionizing radiation-induced senescence). These result in both activation and repression of distinct sets of genes. One prominent consequence is the activation of SASP, which has been argued to occur in all senescent cells ^22, 32, 33^. SASP genes encode a broad repertoire of different proteins, including cytokines, growth factors and matrix metalloproteinases. These affect the physiology of neighboring cells and are also responsible for many of the patho-physiological consequences as documented for example in cancer ^88, 89^. While genes associated with SASP, such as *Cxcl1, Cxcl16* and *IGFBP-6* were slightly upregulated, albeit not significantly (Suppl. Table S1), we did not observe a broad SASP response. *IL6, IL7* and *Ccl2*, prominent members of SASP, were initially downregulated upon loss of Ash2l, while at later time points we observed an increase in *Ccl2* by RT-qPCR (Fig. 5), while these genes were not affected in RNA-seq (Suppl. Table S1). We interpret this as an attempt to activate SASP as part of the senescence response to downregulation of Ash2l but activation of gene expression is inefficient or not possible once H3K4 methylation is lacking. This is also consistent with the finding that MLL1/KMT2A is essential for SASP ^90^.

MAP kinases, including p38MAPK, have been described to be important signaling molecules during senescence and to activate SASP ^91, 92, 93, 94^. Indeed, we observed activation of p38MAPK (Fig. 2). Moreover, the activation of PI3 kinases has been connected to anti-apoptosis in senescent cells as inhibiting this kinase and downstream effectors are senolytic ^27, 28, 95^. Inhibiting MAP kinases by Dasatinib ^96^ and PI3 kinases by Compound 15e, Wortmannin and Quercetin showed senolytic effects in KO2 cells, consistent with these being senescent upon loss of Ash2l.

A number of reports have indicated that gene repression is also associated with senescence. We compared our downregulated genes with those that are associated with senescence in a number of different setups in human primary and cancer cells ^34, 35, 36, 37^. We identified a group of 28 genes that were in common (Fig. 4). GO analysis identified senescence, cell cycle and stress as relevant terms. Several of the genes in this group encode proteins directly linked to senescence. This includes HMGB2, which downregulated early in senescence ^35^, and during aging ^63^. Conversely, the expression of HMGB2 is induced when cell enter the cell cycle form a quiescent state ^64^. A role of HMGB2 in senescence is further supported by its role in inhibiting the spreading of heterochromatin, which ensures expression of SASP genes ^65^. However, the expression of HMGB2 was not sufficient to affect senescence in iMEF cells (Fig. 5).

Moreover, Cenpa, Neil3 and cyclin A2 can interfere with senescence ^51, 97, 98^. In addition, several other member of the M group seem to affect senescence indirectly. Of particular interest is that many of these genes are regulated by FOXM1, as listed above. Indeed, the expression of FOXM1 in iMEF cells slowed the appearance of senescence cells by 2 days (Fig. 5).

Of note is that previous studies have suggested a link of KMT2 complex subunits to senescence. MLL1/KMT2A is frequently translocated in leukemias ^99^. The MLL1-ENL fusion protein, which promotes aberrant proliferation, provokes a DNA damage response and senescence, initially restricting tumor cell proliferation ^100^. The knockdown of SET1A/KMT2F, which is frequently overexpressed in tumors, results in mitotic defects and senescence ^101^. In summary, we identified the trithorax protein Ash2l as a regulator of senescence in fibroblasts. The *Ash2l* KO MEFs displayed many hallmarks of senescent cells but lacked the typical activation of genes encoding SASP factors. Instead, we find that the downregulation of M group genes provide a signature that seems to be relevant for many different senescent phenotypes.

## MATERIALS AND METHODS

### Embryonic fibroblasts

Primary fibroblast cultures from littermate embryos (d.p.c. 13.5) were prepared using mice harboring wild-type or exon 4-floxed alleles of *Ash2l* ^14^ and a single copy of the constitutively expressed transgene CAGGCre-ER™ (strain 004682, The Jackson Laboratory) comprising a fusion between Cre recombinase and the G525R mutant of the mouse estrogen receptor (pMEF, WT: *Ash2l*^+/+^/*Cre-ER* and KO: *Ash2l*^fl/fl^/*Cre-ER*). For the generation of immortalized fibroblast cultures (iMEF) the cells were infected with a retrovirus encoding an shRNA against p19^Arf 23, 24^ and a Blasticidin resistance gene used for the initial selection of infected cells (10 µg/ml Blasticidin, Invivogen, ant-bl). MEF cells were cultivated in DMEM (Thermo Fisher Scientific, 61965059) supplemented with 10% heat-inactivated fetal bovine serum (FBS, Thermo Fisher Scientific, 10270106) and 1% penicillin/streptomycin (Thermo Fisher Scientific, 15070063).

KO2 cells of an early passage were infected with lentiviruses expressing HMGB2- eYFP or FOXM1-eYFP fusion proteins and selected with 20 µg/ml hygromycin B (Goldbio Biotechnology, H-270-1). Pools of cells were then cultured in the presence of hygromycin. The expression of FOXM1-eYFP and HMGB2-eYFP was induced by adding 1 µg/ml Doxycycline (Sigma Aldrich, D9891).

### Induction of recombination

At day 0 cells were treated with 5 nM 4-Hydroxytamoxifen (+HOT) (Tocris, 3412) or with vehicle (-HOT, 100% ethanol). Thereafter cells were passaged when needed to maintain subconfluency or solely supplied with new medium with the addition of HOT or vehicle as before.

### End-point PCR of genomic DNA

Genomic DNA was isolated using the High Pure PCR Template Preparation Kit (Roche, 11796828001) and the PCR performed using the ALLin Red Taq MasterMix (highQu, PCM0201) according to the manufacturers’ instructions. Fragments were analyzed by agarose gel electrophoresis.

### RT-qPCR

Total RNA was isolated with the High Pure RNA Isolation Kit (Roche, 11828665001) and cDNA synthesized with the QuantiTect Reverse Transcription Kit (Qiagen, 205314). A SYBR Green reaction mix (QuantiNova, Qiagen 208054) was employed for the quantitative PCR (qPCR) analyses in a RotorGene 6000 cycler (Corbett/Qiagen). For the individual transcripts, QuantiTect Primer Assays (Qiagen) or primers designed with the Primer3 software (http://primer3.ut.ee) were employed as indicated (see Table 1). The efficiency of all primers used was higher than 95%. The PCR reactions were performed with an initial step at 95 °C for 2 minutes, followed by 40 cycles at 95 °C for 10 s, 60 °C for 10 s, and 72 °C for 5 s and a melting curve analysis. Fold changes in expression were calculated according to Pfaffl ^102^ taking into account the respective primer reaction efficiencies and using *β-glucuronidase* (*Gusb*) as the reference gene as its expression was only little or not affected by the HOT treatment.

**Table 1.**
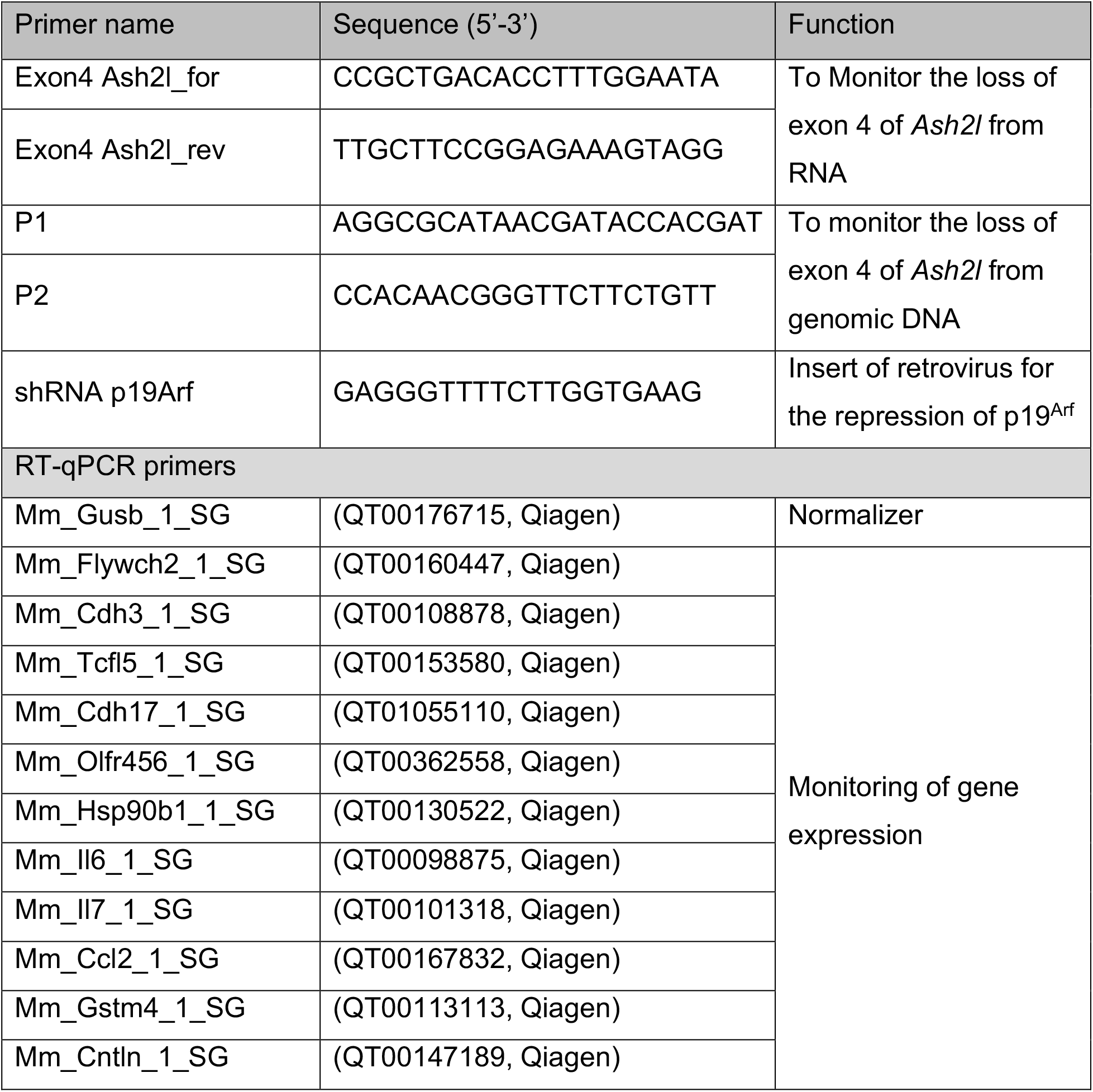
Oligonucleotides

### RNA-seq

From parallel cell cultures the number of cells was determined and RNA purified as noted above. The cell number was used to adjust the amount of ERCC RNA Spike-In Mix (ThermoFisher Scientific, 4456740). Generation of RNA-seq libraries (TruSeq Stranded Total RNA Sample Library Prep, Illumina, 20020596) and next-generation sequencing (NextSeq 500/550 High Output Kit v2.5, 150 cycles, Illumina, 20024907) were performed by the Genomics Core Facility of the Interdisciplinary Center for Clinical Research (IZKF) Aachen of the Faculty of Medicine at RWTH Aachen University.

### Western Blots

Cells were lysed on ice in RIPA buffer (10 mM Tris/HCl, pH 7.4, 150 mM NaCl, 1% Nonidet P-40, 1% deoxycholic acid, 0.1% SDS, protease inhibitor cocktail (1xPIC, Merck, P8340)), containing 10 mM sodium butyrate. After sonication and centrifugation, the protein contents of the supernatants were determined prior to SDS-PAGE and immunoblotting. To verify equal loading the membranes were reversibly stained with 0.2% Ponceau S. The antibodies were applied in 5% low-fat milk, 0.05% Tween20 in PBS. For analyzing p38 phosphorylation, the lysis buffer contained additionally a phosphatase inhibitor cocktail (1 mM ortho-vanadate, 50 nM okadaic acid, 50 nM β-phospho-glycerol, 25 mM NaF, 5 mM EGTA). The antibodies were applied in 5% BSA, 0.05% Tween20 in PBS. The blots were developed with the SuperSignal West Femto Chemiluminescence Substrate (ThermoFisher Scientific, 34094). For details of the antibodies see Table 2.

**Table 2.**
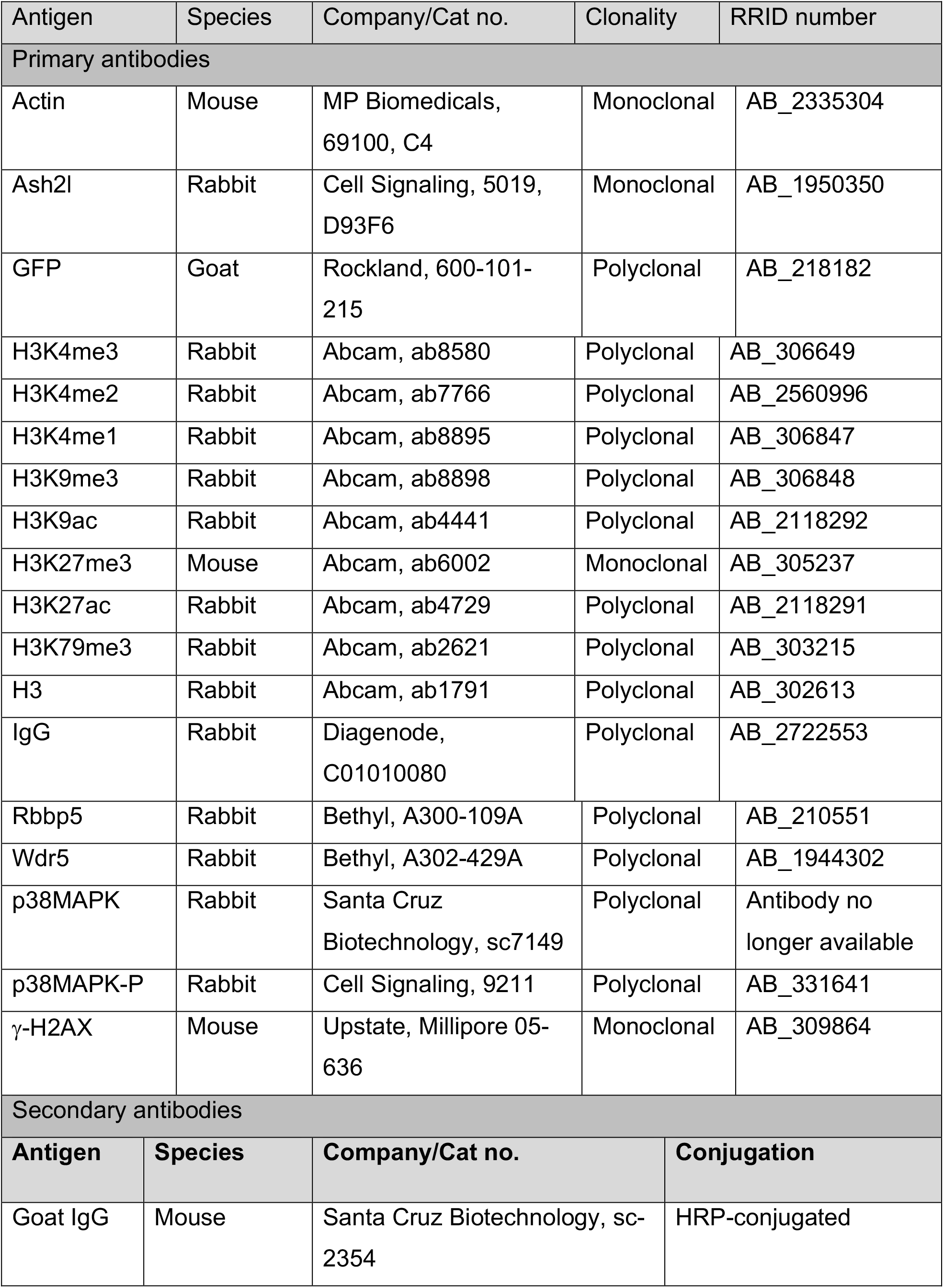
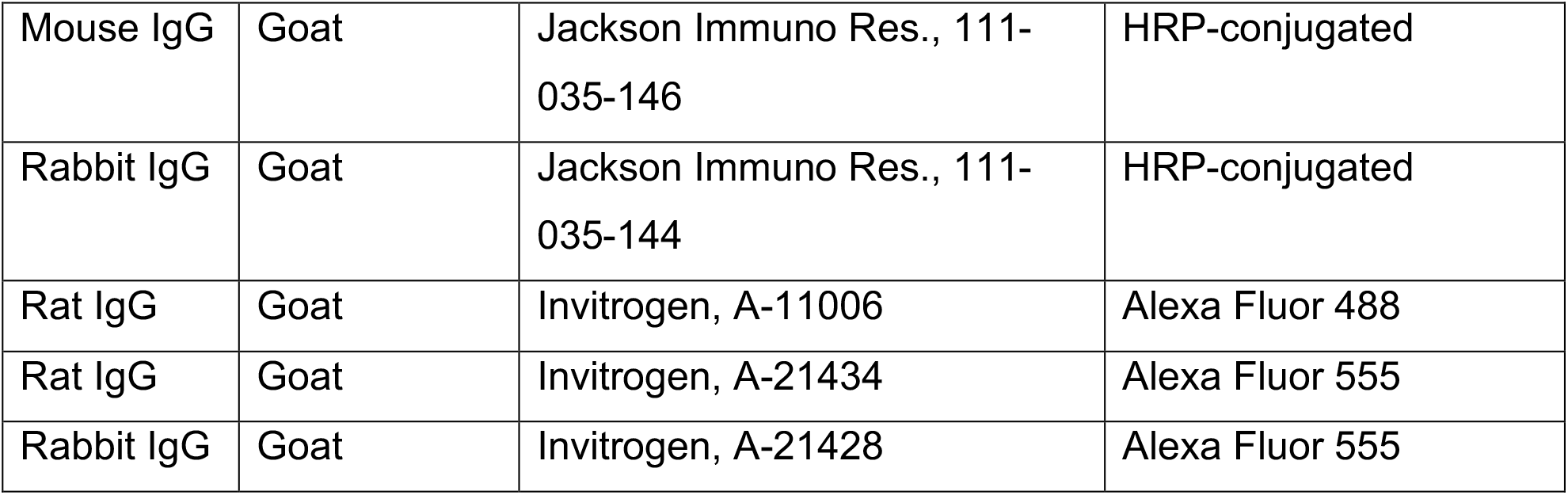
Antibodies

Antibodies
| Antigen | Species | Company/Cat no. | Clonality | RRID number |
| --- | --- | --- | --- | --- |
| Primary antibodies |  |  |  |  |
| Actin | Mouse | MP Biomedicals, 69100, C4 | Monoclonal | AB_2335304 |
| Ash2l | Rabbit | Cell Signaling, 5019, D93F6 | Monoclonal | AB_1950350 |
| GFP | Goat | Rockland, 600-101-215 | Polyclonal | AB_218182 |
| H3K4me3 | Rabbit | Abcam, ab8580 | Polyclonal | AB_306649 |
| H3K4me2 | Rabbit | Abcam, ab7766 | Polyclonal | AB_2560996 |
| H3K4me1 | Rabbit | Abcam, ab8895 | Polyclonal | AB_306847 |
| H3K9me3 | Rabbit | Abcam, ab8898 | Polyclonal | AB_306848 |
| H3K9ac | Rabbit | Abcam, ab4441 | Polyclonal | AB_2118292 |
| H3K27me3 | Mouse | Abcam, ab6002 | Monoclonal | AB_305237 |
| H3K27ac | Rabbit | Abcam, ab4729 | Polyclonal | AB_2118291 |
| H3K79me3 | Rabbit | Abcam, ab2621 | Polyclonal | AB_303215 |
| H3 | Rabbit | Abcam, ab1791 | Polyclonal | AB_302613 |
| IgG | Rabbit | Diagenode, C01010080 | Polyclonal | AB_2722553 |
| Rbbp5 | Rabbit | Bethyl, A300-109A | Polyclonal | AB_210551 |
| Wdr5 | Rabbit | Bethyl, A302-429A | Polyclonal | AB_1944302 |
| p38MAPK | Rabbit | Santa Cruz Biotechnology, sc7149 | Polyclonal | Antibody no longer available |
| p38MAPK-P | Rabbit | Cell Signaling, 9211 | Polyclonal | AB_331641 |
| γ-H2AX | Mouse | Upstate, Millipore 05-636 | Monoclonal | AB_309864 |
| Secondary antibodies |  |  |  |  |
| Antigen | Species | Company/Cat no. | Conjugation |  |
| Goat IgG | Mouse | Santa Cruz Biotechnology, sc-2354 | HRP-conjugated |  |

|  |  |  |  |
| --- | --- | --- | --- |
| Mouse IgG | Goat | Jackson Immuno Res., 111-035-146 | HRP-conjugated |
| Rabbit IgG | Goat | Jackson Immuno Res., 111-035-144 | HRP-conjugated |
| Rat IgG | Goat | Invitrogen, A-11006 | Alexa Fluor 488 |
| Rat IgG | Goat | Invitrogen, A-21434 | Alexa Fluor 555 |
| Rabbit IgG | Goat | Invitrogen, A-21428 | Alexa Fluor 555 |

### *In vitro* SAM assay

Cell lysates were prepared in Co-IP buffer (10 mM HEPES, pH 7.5, 50 mM NaCl, 30 mM Na_4_P_2_O_7_, 50 mM NaF, 0.2% (v/v) Triton X-100, 10% (v/v) glycerol, 5 μM ZnCl_2_, 1xPIC). Lysed cells were mixed vigorously for 45 sec and centrifuged at 16,000 x g and 4°C for 20 min. The protein concentrations of the supernatants were measured and adjusted accordingly. Aliquots were retained for direct Western blot analysis (Input) and otherwise subjected to immunoprecipitation (IP) with protein A-coupled Sepharose beads (Merck, 17-5280-02) pre-incubated with 2 μg anti-RbBP5 antibody or IgG at 4°C for 2 hours. Protein-antibody-bead complexes were washed three times with Co-IP buffer and twice with SAM assay buffer (50 mM Tris-HCl (pH 8.0), 5 mM MgCl_2_, 50 mM NaCl, 1 mM DTT, 1xPIC). Afterwards 2 µg of recombinant human Histone H3.1 (New England BioLabs, M2503), 100 µM S-(5′-adenosyl)-L-methionine chloride dihydrochloride (SAM, Merck, A7007) used as a primary methyl donor molecule, dissolved in 30 µl SAM assay buffer were added to the beads and incubated for 1 h at 31°C with agitation. Subsequently the beads were heated in sample buffer (40 mM Tris pH 6.8, 10 % Glycerol, 2% SDS, bromophenol blue) at 95°C for 10 min and used for SDS-PAGE and immunoblotting.

### Immunoprecipitation (GFP-Trap)

eYFP-tagged fusion proteins were immunoprecipitated using anti-GFP magnetic agarose beads (Chromotek, gtma-20). After induction of gene expression of lentiviral transduced KO2 cells with 1 µg/ml doxycycline, cells were harvested and pelleted. Lysis was performed with ice cold RIPA buffer as described above. A fraction of the lysate was used as input, while the rest was subjected to IP with 5 µl of magnetic beads. IP samples are incubated on an overhead rotator at 4°C for 2 hours and subsequently washed three times in RIPA buffer. The beads were collected in magnetic stand and proteins solubilized in loading buffer as described above.

### Immunofluorescence

Immortalized WT and KO iMEF cells were treated with HOT or vehicle for 5 days, fixed in paraformaldehyde (4% in PBS) at RT for 20 min and permeabilized at RT with TritonX100 (0.2% in PBS) for 5 min. Subsequently, unspecific binding sites were blocked with 20% horse serum (HS) in PBS at 37°C for 30 min. All antibodies were applied in 20% HS/PBS: the primary antibody (specific for Ash2l) at 37°C for 45 min and the secondary antibody (anti-rat Alexa Fluor 488) at 37°C for 30 min. On parallel slides primary antibodies (anti-H3K4me3 or anti-γ-H2AX) were incubated overnight at 4°C followed by the secondary antibody (anti-rat or anti-rabbit Alexa Fluor 555) at 37°C for 30 min. After each incubation the cells were washed 3 times with PBS. Finally, the nuclei were visualized with Hoechst 33258 (Merck, B2883, 2 µg/ml in H2O at RT for 5 min).

### Cell proliferation and SA-β-Galactosidase staining

To measure the proliferation, pMEF and iMEF cells were seeded in triplicates, treated with HOT or vehicle and counted at the days indicated using the CASY® Technology Cell Counter (Innovatis) with three measurements per sample.

Staining for senescence-associated β-galactosidase activity (SA-β-Gal) was performed with cells fixed with glutaraldehyde (0.5% in PBS, RT for 10 min) ^103, 104, 105^. After washing twice with PBS containing 2 mM MgCl_2_(pH 6.0), the staining solution (5 mM K-ferrocyanide, 5 mM K-ferricyanide, 1 mg/ml X-gal in MgCl_2_/PBS, pH 6.0) was applied at 4°C overnight. After washing with H_2_O, the percentage of blue cells vs. the total number of cells was determined.

The senolytics Dasitinib (Cayman Chemical, 11498), Quercetin (Cayman Chemical, 10005169), Compound15e (Enzo Life Science, ALX 270-455) and Wortmannin (Merck, 681675) were dissolved in DMSO and added to cultured cells after 5 days of HOT treatment. Wortmannin was re-added daily.

### Flow cytometry analyses

Cell cycle distribution of iMEF cells was analyzed using flow cytometry using the Foxp3/Transcription Factor Staining Buffer Set (eBioscience, 00-5523) according to the manufacturer’s instructions. After trypsinization, cell pellets were resuspended in 200 µl PBS and 800 µl of Fixation/Permeabilization Buffer under constant vortexing. After incubation at RT for 15 min, 2 ml of Permeabilisation Buffer was added and the cells were centrifuged with 380 x g at RT for 3 min. For cell cycle distribution the fixed cells were resuspended in Hoechst 33258 (Merck, B2883 1 µg/ml H2O). For nocodazole treatment, the iMEF cells were incubated for 7 days with HOT or vehicle and with 100 ng/ml nocodazole (Merck, M1404) in DMSO or with vehicle for 18 h before sample collection. Apoptosis was assessed using the FITC Annexin V Apoptosis Detection Kit and with propidium iodide staining (Immunostep, ANXVKF) according to the manufacturer’s protocol. Measurements were performed using a FACSCanto II system (BD Biosciences) and analyzed with FlowJo software or Flowing Software 2.

### Quantification and statistical analysis

Error bars represent standard deviation (SD) of the mean, unless otherwise indicated. Statistical significance was evaluated by multiple t-test using GraphPadPrism software, unless otherwise indicated.

### Bioinformatics

For RNA-seq, we trimmed sequences using Trim Galore (http://www.bioinformatics.babraham.ac.uk/projects/trim_galore/). The sequences were aligned against the reference genome (UCSC mm9) using STAR for RNA-seq ^106^. We used Gencode/ENSEMBL annotation (Mouse Release M1) to build a gene count matrix with featureCounts ^107^. Next, we used DEseq2 ^108^ to perform differential expression analysis. Genes were normalized only by ERCC (External RNA Controls Consortium) counts. Differentially regulated genes per each cell type and differential promoters are listed in Suppl. Table S1.

GO term/pathway enrichment analyses were performed using the Metascape interface tool (http://metascape.org/gp/index.html) ^109^. For the GO analysis in Figure 4 we employed g:Profiler ^110^.

Sequencing data of the RNA-seq experiments have been deposited in Gene Expression Omnibus (Suppl. Table S1: https://www.ncbi.nlm.nih.gov/geo/query/acc.cgi, accession number GSE165458, password izyrokiwhropdwz).

## ACKNOWLEDGMENTS

We thank Dr. M. Eilers (University of Würzburg, Germany) for providing the siRNA expression construct that targets the *p19*^*ARF*^ mRNA, J. Stahl for expert technical assistance and L. Gan und J. Hübner of the Genomics Facility of the Interdisciplinary Center for Clinical Research (IZKF) Aachen, Faculty of Medicine, RWTH Aachen University for support in the gene expression analysis. Simulations were performed with computing resources granted by RWTH Aachen University under project rwth0751 (to M.B.). The work was funded in part by grants from the German Research Foundation DFG (LU466/17-1 and 17-2 to B.L.) and the Interdisciplinary Center for Clinical Research (IZKF) Aachen (to I.C.).

## AUTHOR CONTRIBUTIONS

A.B., A.T.S., R.S.B., W.L., P.B. and J.L.F. performed the wet experiments; R.S.B., C.C.K., M.B. and I.C. carried out bioinformatics analyses; J.L.F and B.L. developed the project; A.B., R.S.B., C.C.K., M.B., I.C., J.L.F and B.L. analyzed data; J.L.F, I.C. and

B.L. wrote the initial version of the paper; all authors read and approved the final manuscript.

## ADDITIONAL INFORMATION

## Supplementary information

4 Figures, 2 Tables

### Ethical approval and informed consent

All procedures were approved by the Institute for Laboratory Animal Science and the local government (LANUV NRW, AZ 84-02.04.2013.A182) in accordance with German and EU regulations (also indicated in Material and Methods section).

