## Supplemental tables and figures for "Induction of senescence upon loss of the Ash2l core subunit of H3K4 methyltransferase complexes"

Mirna Barsoum: 0000-0003-0838-8137

Ivan G. Costa: 0000-0003-2890-8697

Juliane Lüscher-Firzlaff: 0000-0001-9265-8536

Bernhard Lüscher: 0000-0002-9622-8709

**Suppl. Table S1**

RNA-seq data: https://www.ncbi.nlm.nih.gov/geo/query/acc.cgi, accession number GSE165458, password izyrokiwhropdwz

**Suppl. Table S2**

| | | Expression change between (Log2 fold)* | | | |
|---|---|---|---|---|---|
| **Protein** | **Protein coding gene** | **KO2+/KO2-** | **Significance**** | **KO1+/KO1-** | **Significance** |
| **Cyclines and Cyclin-dependent kinases** | | | | | |
| Cyclin A2 | *Ccna2* | -1.31 | + | -1.29 | + |
| Cyclin B1 | *Ccnb1* | -1.12 | + | -1.21 | + |
| Cyclin B2 | *Ccnb2* | -1.85 | + | -1.91 | + |
| Cyclin D2 | *Ccnd2* | -1.32 | + | -0.42 | |
| Cyclin E2 | *Ccne2* | -1.28 | + | -1.04 | |
| Cdk1 | *Cdk1* | -1.34 | + | -1.19 | + |
| **Transcription factors and cofactors involved in cell cycle regulation** | | | | | |
| E2f1 | *E2f1* | -1.54 | + | -0.96 | |
| E2f2 | *E2f2* | -1.65 | + | -1.41 | + |
| E2f8 | *E2f8* | -1.36 | + | -0.91 | |
| Rbp1 | *Rbp1* | -3.38 | + | -3.96 | + |
| p103 | *Rbl1* | -1.36 | + | -1.12 | |
| **Replication** | | | | | |
| Cdc6 | *Cdc6* | -1.22 | + | -0.94 | |
| Mcm2 | *Mcm2* | -1.40 | + | -0.83 | |
| Mcm3 | *Mcm3* | -1.31 | + | -0.78 | |
| Mcm4 | *Mcm4* | -1.16 | + | -0.81 | |
| Mcm6 | *Mcm6* | -1.71 | + | -1.09 | |
| Mcm10 | *Mcm10* | -1.21 | + | -0.83 | |
| Geminin | *Gmnn* | -1.41 | + | -1.05 | |

* Expression analysis of genes associated with the cell cycle and with replication (excerpted from the RNA-seq data (complete list of genes in Table S1))
** Described in the main manuscript.

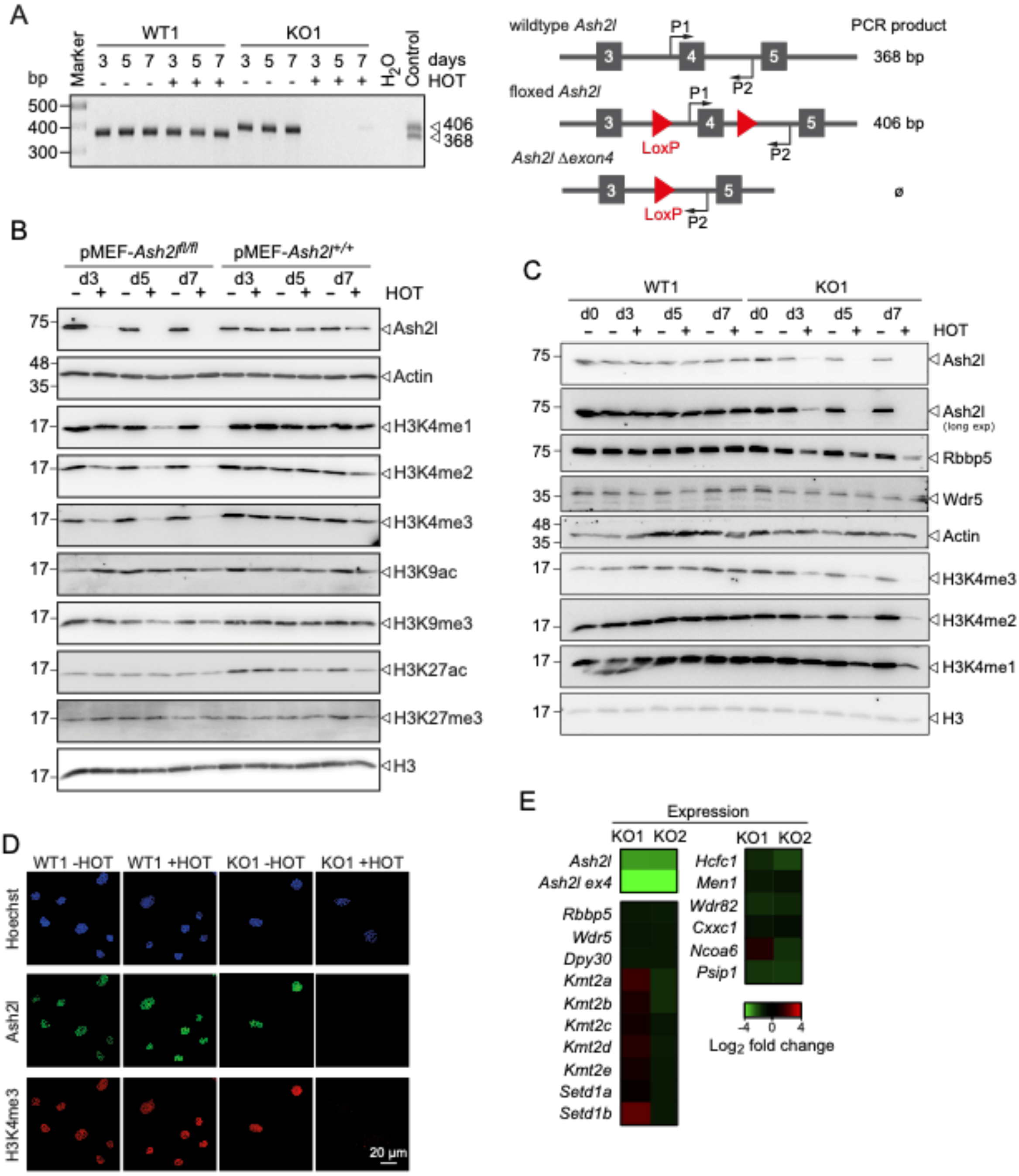


Suppl. Figure S1 (related to Fig. 1)

A. Wild type (WT1) and *Ash2l*$^{fl/fl}$ (KO1) immortalized fibroblasts (iMEF1) were treated with 4-hydroxy tamoxifen (HOT, 5 nM) or vehicle for the times indicated. Genomic PCR with primers P1 and P2 identifies a 368 bp fragment in WT cells and a 406 bp fragment in the KO cells. Recombination deletes the target sequence of P1.

B. *Ash2l*$^{wt/wt}$:*Cre-ER*$^{TM2}$ and *Ash2l*$^{fl/fl}$:*Cre-ER*$^{TM2}$ primary embryonal fibroblasts (pMEFs) were treated ± HOT for 3, 5 or 7 days. The cells were lysed and the indicated proteins analyzed by Western blotting.

C. *Ash2l*$^{wt/wt}$:*Cre-ER*$^{TM2}$ (WT1) and *Ash2l*$^{fl/fl}$:*Cre-ER*$^{TM2}$ (KO1) immortalized fibroblasts (iMEF1) were treated ± HOT for 0, 3, 5 or 7 days. The cells were lysed and the indicated proteins analyzed by Western blotting.

D. WT1 and KO1 cells were treated ± HOT for 5 days. The cells were fixed and the DNA stained using Hoechst. Ash2l (green) and H3K4me3 (red) were labeled using specific antibodies.

E. The expression of RNA encoding subunits of KMT2 complexes (left) and associated proteins (right) was analyzed from the RNA-seq experiments in KO1 and KO2 cells (Table S1). Compared are the signals after HOT treatment for 5 days compared to vehicle treated controls.

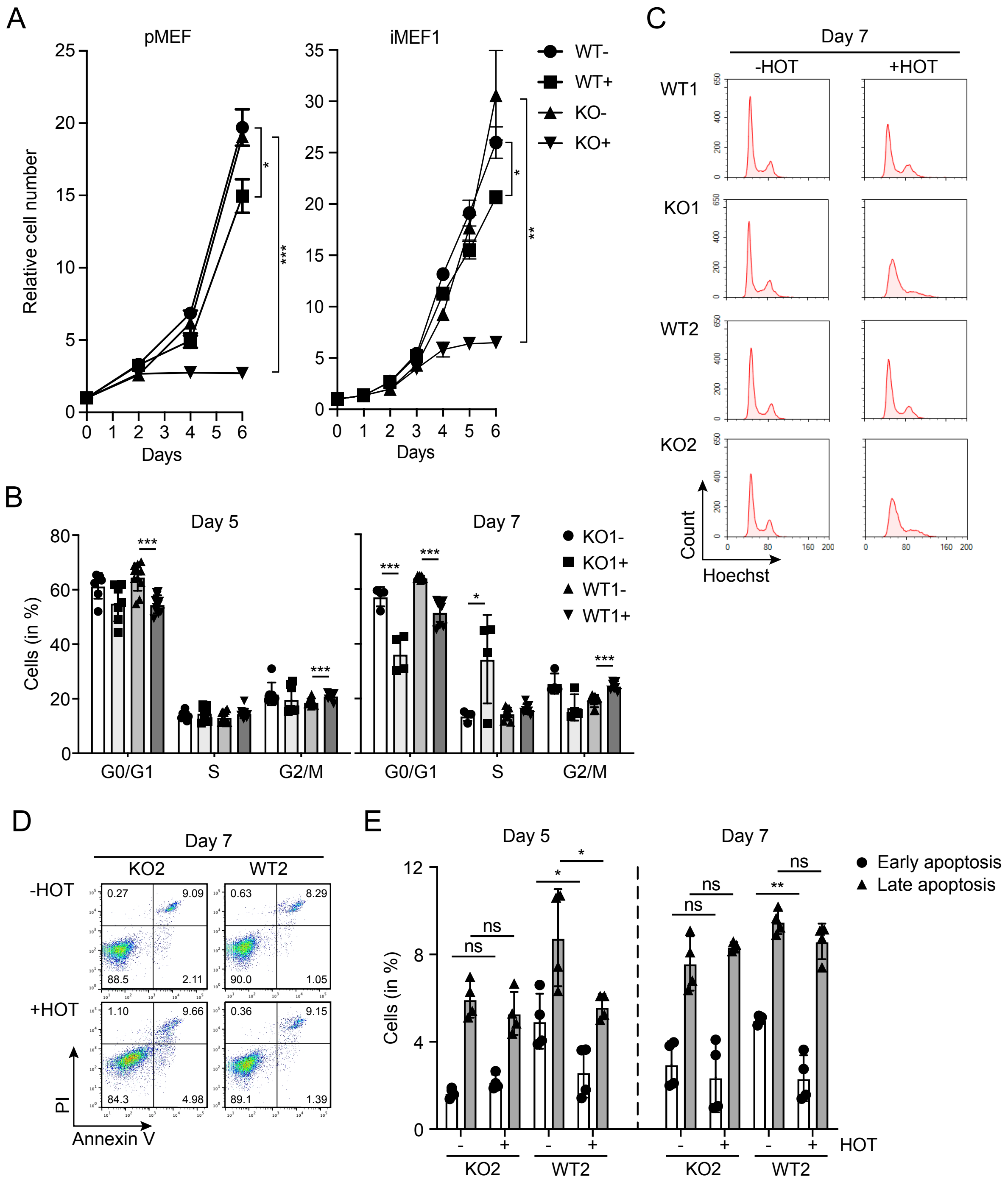


Suppl. Figure S2 (related to Fig. 1)

A. Primary fibroblasts (pMEF) and iMEF1 cells were treated ± HOT (5 nM) and counted as indicated. Mean values ± SD (n = 3, with 2 measurements for each time point. Statistical analyses refers to day 6: * <0.05, ** < 0.01, *** < 0.001).

B. Cell cycle analysis using flow cytometry of fixed and Hoechst stained WT1 and KO2 cells treated ± HOT for 5 or 7 days. Mean values ± SD (n = 6-10; ** < 0.01, *** < 0.001).

C. Exemplary flow cytometry analysis of WT1, KO1, WT2 and KO2 cells treated ± HOT for 7 days. Cells were fixed and the DNA stained using Hoechst.

D. Exemplary flow cytometry analysis of WT2 and KO2 cells treated ± HOT for 7 days. Cells were stained for Annexin V and with propidium iodide.

E. Analysis as in panel D. Early apoptosis refers to Annexin V positive cells, late apoptosis to Annexin V and propidium iodide double positive cells. Mean values ± SD (n = 4; * <0.05, ** p<0.01; ns, not significant).

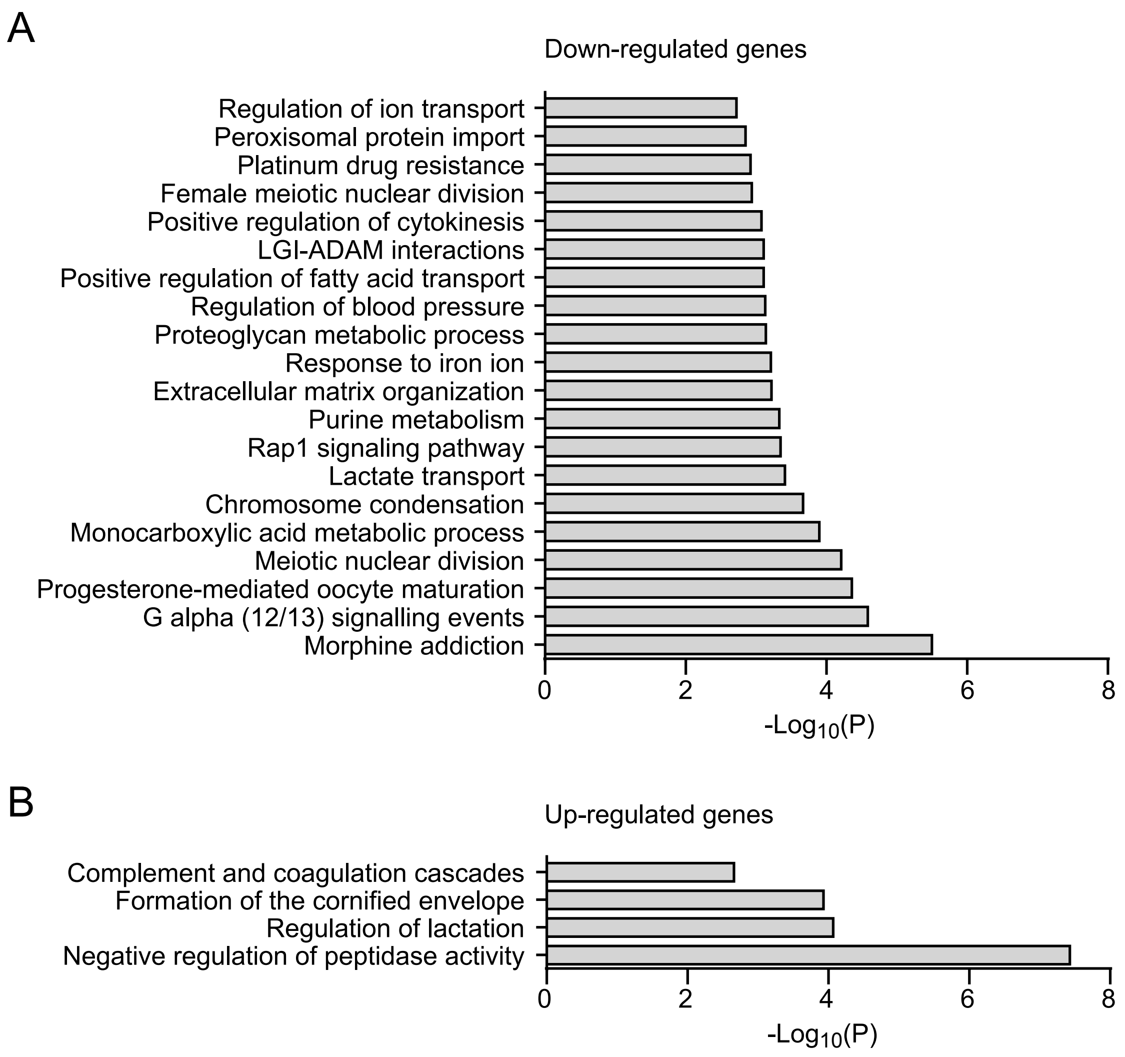


Suppl. Figure S3 (related to Figs. 3 and 4)

A. GO analysis of down-regulated genes in response to HOT treatment.

B. GO analysis of up-regulated genes in response to HOT treatment.

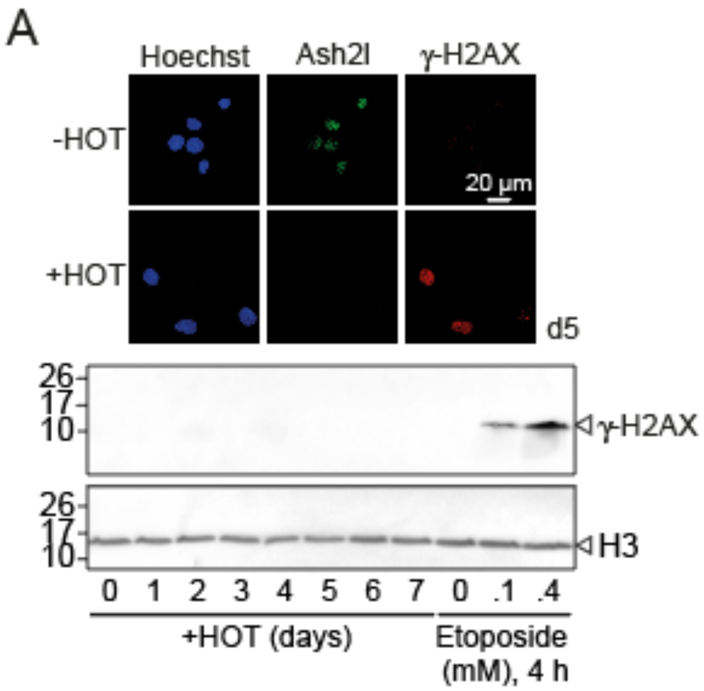


Suppl. Figure S4 (related to Fig. 5)

A. KO1 cells were treated ± HOT for 5 days. The cells were fixed and the DNA stained (Hoechst). Ash2l and γ-H2AX were visualized using specific antibodies (upper panel). KO2 cells were treated ± HOT for the indicated days. For control untreated KO2 cells were incubated with etoposide for 4 hrs. RIPA cell lysates were prepared and H3 and γ-H2AX were detected on Western blots.

B. KO2 cells were infected with lentiviruses expressing HMGB2-eYFP. Pools of cells were treated with doxycycline (Dox). The expression of HMGB2-eYFP was analyzed using Western blotting (WB). Input refers to 2.5% of total cell lysate, the remainder was used for immunoprecipitating the fusion protein (IP). Prior to specific protein detection, the membranes were stained with Ponceau S to verify equal loading of input lysates.
